# A single dose of replication-competent VSV-vectored vaccine expressing SARS-CoV-2 S1 protects against virus replication in a hamster model of severe COVID-19

**DOI:** 10.1101/2021.01.29.428442

**Authors:** Delphine C. Malherbe, Drishya Kurup, Christoph Wirblich, Adam J. Ronk, Chad Mire, Natalia Kuzmina, Noor Shaik, Sivakumar Periasamy, Matthew A. Hyde, Julie M. Williams, Pei-Yong Shi, Matthias J. Schnell, Alexander Bukreyev

**Affiliations:** Department of Pathology, University of Texas Medical Branch, Galveston, TX 77555, U.S.A.; Galveston National Laboratory, Galveston, TX 77555, U.S.A.; Department of Microbiology and Immunology, Jefferson University, Philadelphia, PA 19107, U.S.A.; Jefferson Vaccine Center, Thomas Jefferson University, Philadelphia PA 19107, U.S.A.; Department of Biochemistry and Molecular Biology, University of Texas Medical Branch, Galveston, TX 77555, U.S.A.; Institute for Human Infections and Immunity, University of Texas Medical Branch, Galveston, TX 77555, U.S.A.; Department of Microbiology & Immunology, University of Texas Medical Branch, Galveston, TX 77555, U.S.A.

**Keywords:** COVID-19, SARS-CoV-2, vaccine, hamster, VSV vector

## Abstract

The development of effective countermeasures against Severe Acute Respiratory Syndrome Coronavirus 2 (SARS-CoV-2), the agent responsible for the COVID-19 pandemic, is a priority. We designed and produced ConVac, a replication-competent vesicular stomatitis virus (VSV) vaccine vector that expresses the S1 subunit of SARS-CoV-2 spike protein. We used golden Syrian hamsters as animal model of severe COVID-19 to test the efficacy of the ConVac vaccine. A single vaccine dose elicited high levels of SARS-CoV-2 specific binding and neutralizing antibodies; following intranasal challenge with SARS-CoV-2, animals were protected from weight loss and viral replication in the lungs. No enhanced pathology was observed in vaccinated animals upon challenge, but some inflammation was still detected. The data indicate rapid control of SARS-CoV-2 replication by the S1-based VSV-vectored SARS-CoV-2 ConVac vaccine.

## INTRODUCTION

The current pandemic of coronavirus induced disease 2019 (COVID-19) caused by severe acute respiratory syndrome coronavirus 2 (SARS-CoV-2) has resulted in 36.4 million confirmed cases of human infections worldwide, including over two million confirmed deaths (as of January 17, 2021) (WHO, 2021). There are currently no approved vaccines or therapeutics against SARS-CoV-2. Multiple vaccines are being developed, including inactivated virus, viral vector-based vaccines, protein subunits and mRNA vaccines (reviewed by Krammer (Krammer, 2020)). The main antigen of these vaccines is the full-length spike (S) protein of SARS-CoV-2, since the virus uses its S protein to mediate cell entry.

Non-segmented negative strand RNA viruses including vesicular stomatitis virus (VSV), which replicates systemically, represent highly potent vaccine platforms (Bukreyev et al., 2006). These vaccine platforms are capable of stably expressing foreign open reading frames flanked with transcriptional gene-start and gene-end signals specific for the viral vector’s polymerase. A number of vaccine candidates for Middle East respiratory syndrome coronavirus (MERS-CoV), SARS-CoV, SARS-CoV-2 and porcine coronavirus utilized VSV and rabies virus as vector backbones to express the spike protein as immunogen (Faber et al., 2005; Kapadia et al., 2008; Kato et al., 2019; Ke et al., 2019; Wirblich et al., 2017). We previously developed a MERS vaccine candidate in which the S1 domain of the MERS spike protein was fused to the C-terminal part of the rabies virus glycoprotein to allow incorporation and display on the surface of virions (Wirblich et al., 2017). We reasoned that removal of the S2 domain should steer the immune response towards the receptor-binding domain which is located in the S1 domain and is the target of neutralizing antibodies. To expedite COVID-19 vaccine development, we took this proven strategy to generate a SARS-CoV-2 vaccine candidate in which the S2 domain was replaced with the C-terminal part of the VSV glycoprotein as a membrane anchor and expressed in a replication-competent VSV vector.

We assessed the immunogenicity and efficacy of our vaccine in the recently established golden Syrian hamster model of COVID-19 (Chan et al., 2020; Imai et al., 2020; Sia et al., 2020). Golden Syrian hamsters are susceptible to intranasal infection with wild type SARS-CoV-2 without the need for virus adaptation, and develop severe clinical manifestations similar to those observed in human COVID-19 patients with a severe disease (Chan et al., 2020; Imai et al., 2020; Sia et al., 2020).

## RESULTS

### Design and development of the VSV-SARS-CoV-2 vaccine ConVac

To design our ConVac vaccine construct, we decided to use only the S1 domain of SARS-CoV-2 spike S for several reasons. First, the S1 region contains the necessary neutralizing epitopes with the receptor-binding domain being most important (Ho, 2020). Second, both prefusion and postfusion S2 structures present potential drawbacks as vaccine constructs, and vaccines expressing the full length S spike protein may generate both prefusion and postfusion spikes as observed in *in vitro* mammalian cell culture production systems (Cai et al., 2020). Because of its high level of glycosylation it has been proposed that the S2 postfusion configuration may act as an immunological decoy by eliciting immunodominant non-neutralizing responses while stabilization of prefusion S2 may affect bonds between protomers thus affecting the overall trimer stability (Cai et al., 2020). The third reason for the use of the S1 only approach is that the S2 domain is more conserved between different coronaviruses (Huang et al., 2020; Walls et al., 2020), and we hypothesized that in previously coronavirus-infected humans the immune response might predominantly target S2 rather than S1. Lastly, previous research with a rabies virus-vectored vaccine indicated that the expression of the full-length coronavirus S protein interfered with the transport of the rabies virus G protein and reduced vaccine viral titers dramatically (Wirblich et al., 2017). Moreover, VSV G-deleted viruses grow to a lower titer of about 10^7^ PFU, whereas VSV G-containing viruses can reach titers up to 1-5×10^8^, making vaccine production more efficient (Kretzschmar et al., 1997; Schnell et al., 1996). These reasons led us to develop a VSV-based vaccine construct containing only the membrane-anchored S1 domain by substituting the S2 domain of the SARS-CoV-2 spike protein with the C-terminal region of the VSV glycoprotein.

Thus, following the strategy we previously employed to generate a membrane-anchored S1 domain of MERS (Wirblich et al., 2017) we replaced the S2 domain of SARS-CoV-2 (aa 682-1284) with the C-terminal 70 amino acids of VSV G (Fig. 1A). This portion contains the complete cytoplasmic tail of the VSV glycoprotein, as well as the transmembrane domain and a short membrane-proximal portion of the ectodomain of VSV-G. The VSV-G tail serves as a membrane anchor for S1 and allows incorporation into VSV particles. We recovered the virus as described previously (Kurup et al., 2015) and performed immunofluorescence staining with antibodies directed against S1 or VSV glycoprotein (Fig. 1B). Vero cells infected with ConVac expressed both S1 and VSV glycoprotein. No S1 signal was detected in cells infected with a control virus expressing Hendra virus G. We also performed western blotting to assess protein expression (Fig. 1C). Using polyclonal antiserum directed against the S1 domain we detected a protein of 160 kDa and a shorter band of approximately 100 kDa. The smaller band presumably represents unglycosylated or partially glycosylated protein, whereas the larger band represents the fully glycosylated mature protein. VSV glycoprotein expression was noticeably reduced compared to the control virus expressing GFP. To assess the impact on viral replication, we evaluated growth kinetics in Vero cells (Fig. 1D). ConVac grew to titers 2-4×10^8^ FFU/ml, which are about 5-fold lower than the control virus expressing GFP and modestly lower compared to a recombinant VSV expressing Hendra virus G. Similar titers were obtained in human lung and baby hamster kidney cells (data not shown).

**Figure 1.**
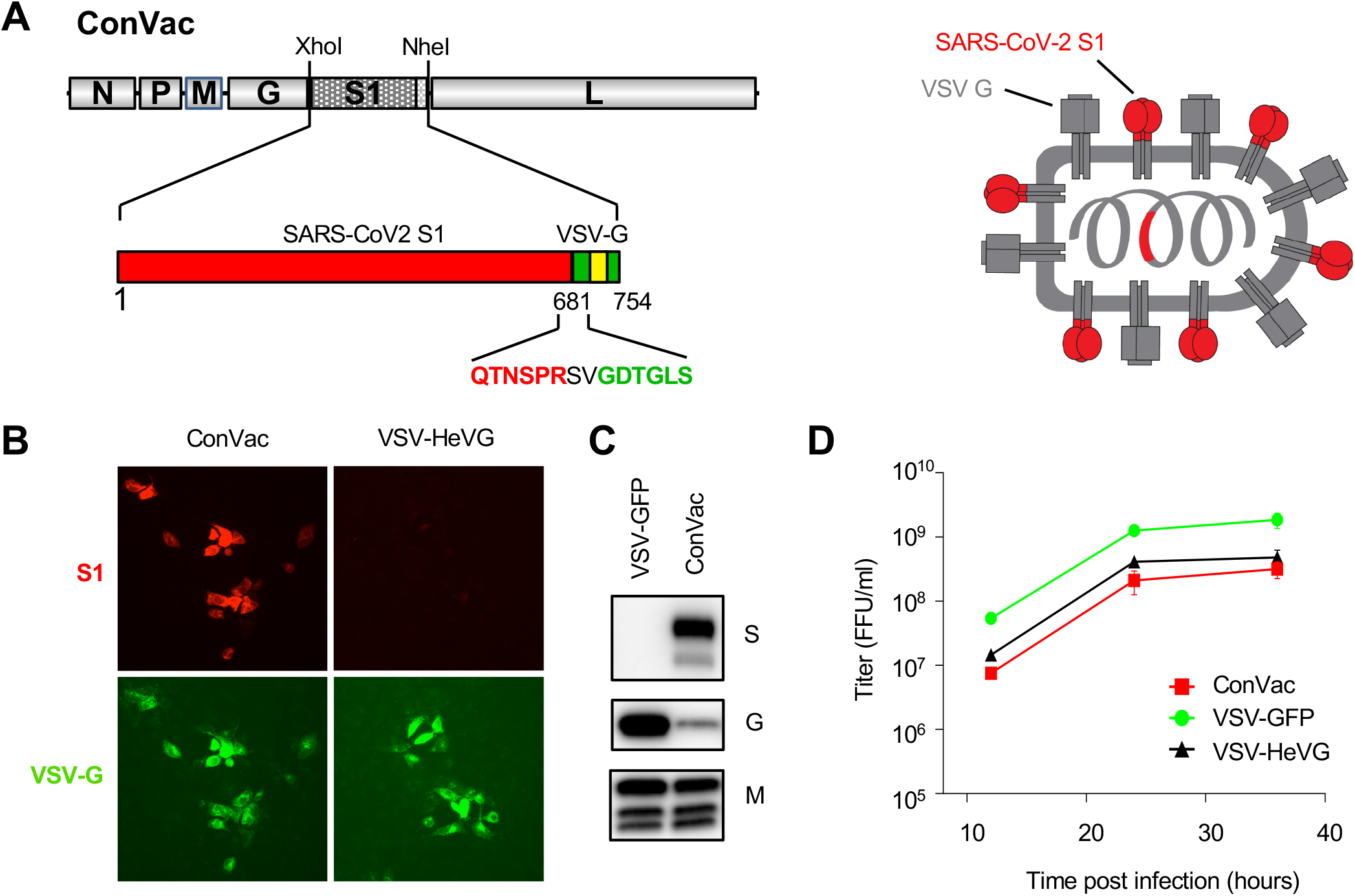
Generation and characterization of ConVac. (A) Left: genome structure of the VSV vector expressing the membrane anchored SARS-CoV2 S1 domain. The S1 domain of SARS-CoV2 spike protein (aa 1-681) was joined to the C-terminal 70 amino acids of the VSV glycoprotein. The fusion construct was inserted between the glycoprotein and polymerase genes of VSV. The S1 domain is shown in red and the VSV G tail in green. The transmembrane domain of the VSV glycoprotein is indicated by a yellow box. The amino acids at the junction between S1 and VSV glycoprotein are highlighted. Right: schematic representation of the vaccine construct which shows the two transmembrane proteins anchored in the membrane. (B) Immunofluorescence staining of Vero E6 cells infected with ConVac and a control virus expressing Hendra virus G (VSV-HeVG). The cells were fixed and permeabilized 10 hours after infection and stained with fluorescently labeled monoclonal antibody CR3022 against the S1 domain (shown in red) and two monoclonal antibodies against the VSV glycoprotein (shown in green). (C) Western blot analysis of BSR cells infected with Convac and a control VSV virus expressing GFP. Protein lysates were resolved on 4-20% polyacrylamide gradient gels and transferred to nitrocellulose membranes. The membranes were probed with polyclonal antiserum against the S1 domain (upper panel), monoclonal antibodies against the VSV glycoprotein (middle panel) and a monoclonal antibody against the VSV matrix protein (lower panel). VSV glycoprotein expression was significantly reduced in cells infected with ConVac whereas matrix protein expression was only modestly affected. (D) Viral growth curve on Vero cells. The cells were infected at an MOI of 0.05 PFU with ConVac, or VSV expressing GFP or another control virus expressing Hendra virus glycoprotein. Supernatants were collected 12, 24, and 36 hours post infection and titrated on Vero E6 cells.

### Development of the hamster model of COVID-19

To determine if golden Syrian hamsters are a suitable animal model to study the protective efficacy of the vaccine, eight female golden Syrian hamsters were infected intranasally with 10^5^ PFU of SARS-CoV-2 strain USA_WA1/202 (Harcourt et al., 2020) at day 0 in a volume of 100 μl, and four female hamsters were mock-infected with 100 μl of phosphate buffered saline. Animals were monitored daily for weight loss (Fig. 2A). Viral load determination in the lungs (Fig. 2B) and nasal turbinates (Fig. 2C) was performed 2 and 4 days post challenge. At day 4, SARS-CoV-2 infected hamsters had lost an average of 7.3% of their initial body weight, while mock-infected animals did not display any weight loss (Fig.2A). Viral loads were both higher in the lungs (mean viral load of 3.0 x 10^6^ PFU/g) and in the nasal turbinates (2.7 x 10^6^ PFU/g) at day 2 compared to day 4 (3.4 x 10^5^ PFU/g of lung and 4.1 x 10^4^ PFU/g nasal turbinate) (Fig. 2B,C). Lung histopathological changes were assessed in two animals of each group at each time point (criteria displayed in Supplemental Table 1 (Matute-Bello et al., 2011)). The SARS-CoV-2 infected hamsters displayed lung pathology consistent with typical interstitial pneumonia at both time points (Fig. 2D) while some alveolar changes and septal thickening (Fig. 2I) were also observed in the mock-infected animals that were a consequence of the CO_2_ euthanasia method used in this pilot study. In SARS-CoV-2 infected hamsters, widespread inflammatory changes consisting of small to large inflammatory foci were observed (Fig. 2E,F). In addition, infiltration of airways with inflammatory cells (Fig. 2G,J) and a moderate level of inter-alveolar septal thickening were noted in SARS-CoV-2 infected animals (Fig. 2H,J). Thus, lung pathology observed in our pilot study indicates that the golden Syrian hamster is a suitable animal model reproducing the typical interstitial pneumonia caused by SARS-CoV-2 in humans. In addition, our findings are in agreement with others as while this manuscript was in preparation, other teams reported that SARS-CoV-2 infection of hamsters causes a severe lung disease, with viral replication in the upper and lower respiratory tract (Chan et al., 2020; Imai et al., 2020; Sia et al., 2020).

**Figure 2.**
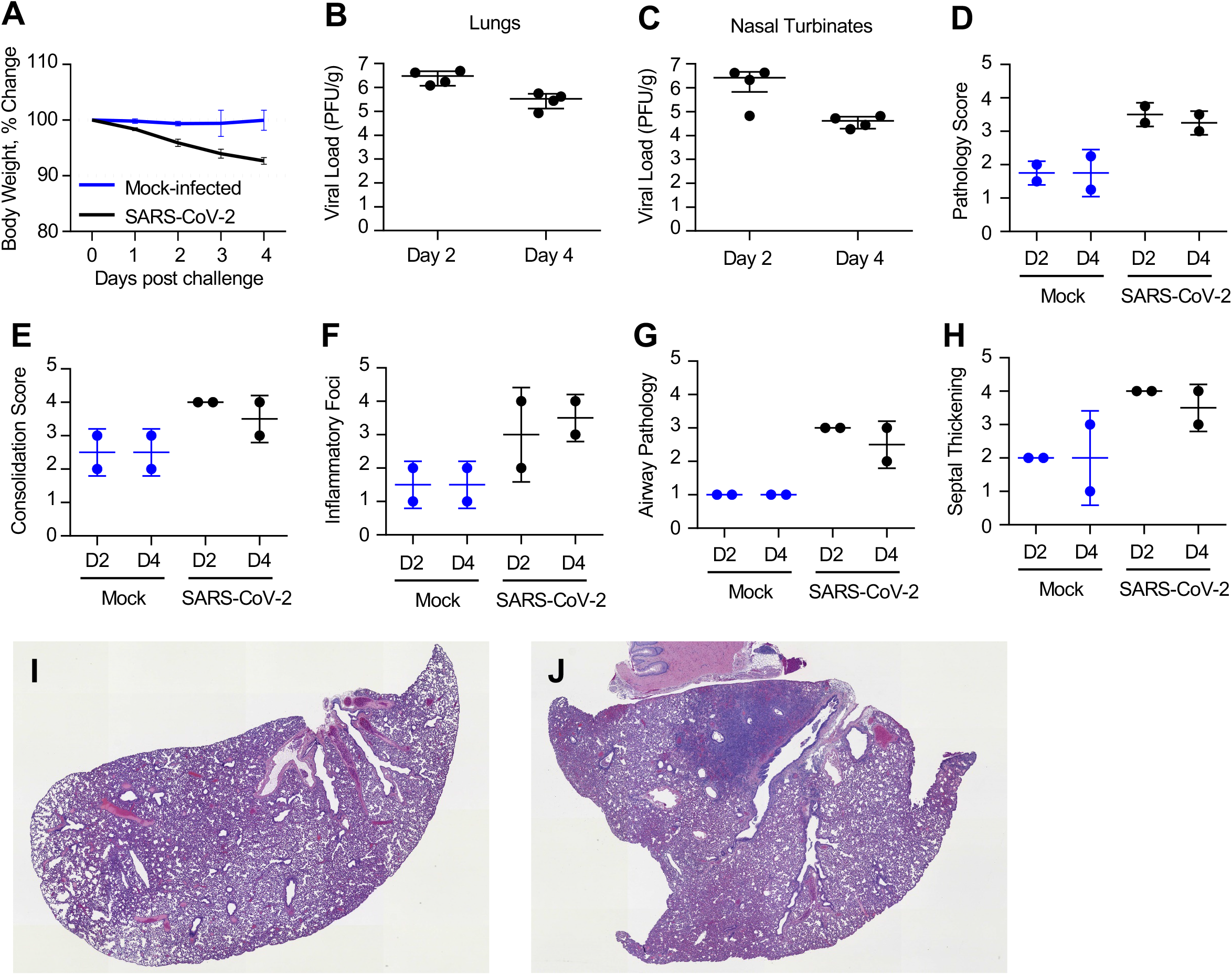
Pathogenicity and replication of SARS-CoV-2 in golden Syrian hamsters. Hamsters were challenged intranasally with 10^5^ PFU SARS-CoV-2 at day 0 and monitored for 4 days. (A) % change in body weight. SARS-CoV-2 group is shown in black, and mock infected group in blue. Viral loads were determined in lung (B) and nasal turbinate (C) tissues 2 days and 4 days post challenge. Right lungs and nasal turbinates from each animal were homogenized in media and titrated on Vero E6 cells. No virus was detected in the mock infected animals (data not shown). N = 4 for mock control group (2 hamsters euthanized at each timepoint) and N = 8 for VSV vaccine group (4 hamsters euthanized at each timepoint). Comparative pathology scores for lungs from SARS-CoV-2-infected and mock-infected hamsters were determined 2 days (D2) and 4 days (D4) post challenge (D-H). Scores for overall lung pathology (D), and individual criteria including consolidation or extent of inflammation (E), type inflammatory foci (F), airway pathology (G) and septal thickening (H) are displayed. The pathology scores (mean) were calculated based on the criteria described in Supplemental Table 1. Data represent mean ± SD, N = 2 for each group at each timepoint. (I) Day 4 mock-infected lung displaying minimal pathologic changes in airways. Note a septal thickening and alveolar wall damage (likely due to CO_2_ euthanasia). (J) Day 4 SARS-CoV2 infected lung displaying widespread inflammatory change, septal thickening and airway infiltration.

### ConVac vaccine induces a robust antibody response in hamsters

The immunogenicity and efficacy of the ConVac vaccine was evaluated in the golden Syrian hamster animal model. Hamsters at 12 animals per group were vaccinated once intramuscularly on day 0 (Fig. 3A) with 2×10^7^ FFU/animal of ConVac, while the control group remained naive. On day 10 post vaccination, one vaccinated hamster was euthanized due to hind leg paralysis, and on day 11 another hamster was found dead after being lethargic, scruffy, hunched and with heavy breathing the day before. All other animals demonstrated no disease symptoms. At day 31, vaccinated and control hamsters were challenged intranasally with a dose of 10^5^ PFU of the SARS-CoV-2 isolate USA_WA1/202 (Harcourt et al., 2020). Serum samples were assayed longitudinally to determine SARS-CoV-2 S1 specific binding antibody responses (Fig. 3). Testing of the binding to purified S1 protein demonstrated induction of IgG EC_50_ titers ranging from 505 to 5,096 with a mean titer of 2,279 (Fig. 3B) after vaccination. One control animal demonstrated presence of SARS-CoV-2 antibodies by ELISA but was negative for SARS-CoV-2 neutralizing antibodies. The S1-specific Th1 isotype response IgG2/3 titers were also determined (Fig. 3C); the ConVac vaccine elicited titers ranging from 39 to 285 with a mean titer of 139, mirroring the total IgG responses albeit at a lower level. The SARS-CoV-2 challenge did not elicit an anamnestic binding antibody response in the ConVac-vaccinated animals, as the binding antibody titers did not increase between the time points before and after the challenge (day 28 vs day 34 and day 28 vs day 46, P>0.05 for both IgG and IgG2/3, Wilcoxon test). Neutralization of SARS-CoV-2 (Fig. 3D) was assessed with the recombinant SARS-CoV-2 expressing Neon Green protein (SARS-CoV-2-mNG) using the fluorescent signal as a readout for viral replication thus enabling a fast and quantitative evaluation (Xie et al., 2020). The ConVac vaccine elicited 50% SARS-CoV-2 neutralizing titers ranging from 1:85 to 1:886 with a mean titer of 1:369 (Fig. 3E). Taken together, our data show that a single vaccine dose induced a robust SARS-CoV-2-specific antibody response.

**Figure 3.**
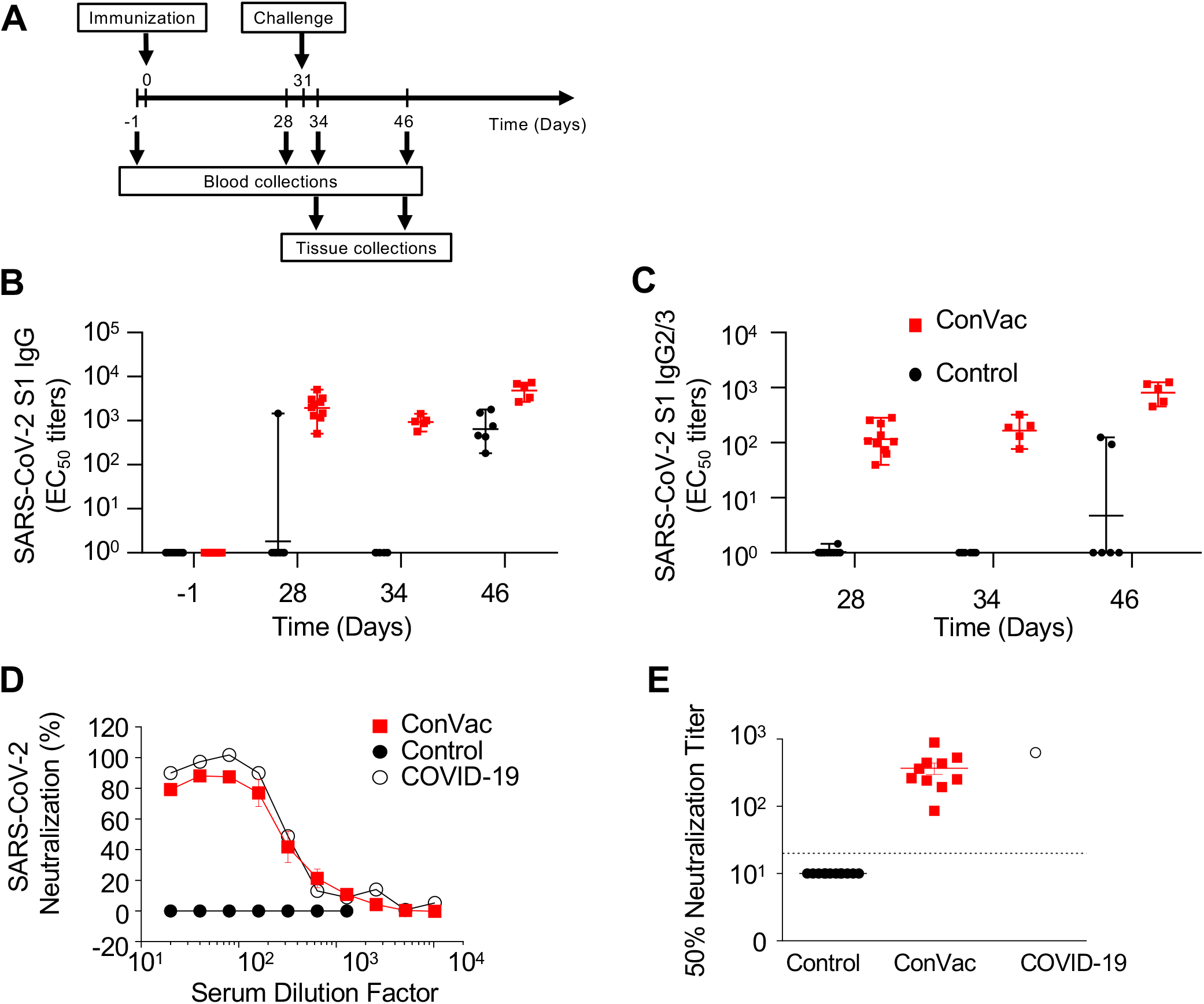
ConVac elicits SARS-CoV-2 S1-specific binding and neutralizing antibodies. The study schedule (A) indicates the immunization, challenge and collection time points. Sera collected during the immunogenicity phase (days −1, and 28) and the challenge phase (day 34 is 3 days post challenge and day 46 is 15 days post challenge) were assessed for their ability to bind to SARS-CoV-2 spike protein S1. Total IgG (B) and IgG2/3 (C) responses are displayed. Sera collected 28 days post inoculation were assessed for their ability to neutralize SARS-CoV-2. Percent neutralization (D) and 50% neutralization titers (E) are displayed. The limit of detection is indicated by the dotted line. A plasma sample from a COVID-19 convalescent human subject was included as positive control (empty circle). The ConVac group is shown in red, the control group in black.

### ConVac vaccine significantly decreases SARS-CoV-2 viral load in the lungs

After the challenge, the hamsters were monitored for 15 days. Animals were checked daily for body weight and clinical signs of disease. In the control group, a significant loss of weight compared to the vaccine group (P=0.0002) was detected, and at the peak of weight loss, which occurred at day 5, the average weight loss was 9.1%, (Fig. 4A). None of the control or the vaccinated animals reached moribund state, but several control animals had higher disease scores due to a weight loss greater than 10% of their initial body weight (Fig. 4B).

**Figure 4.**
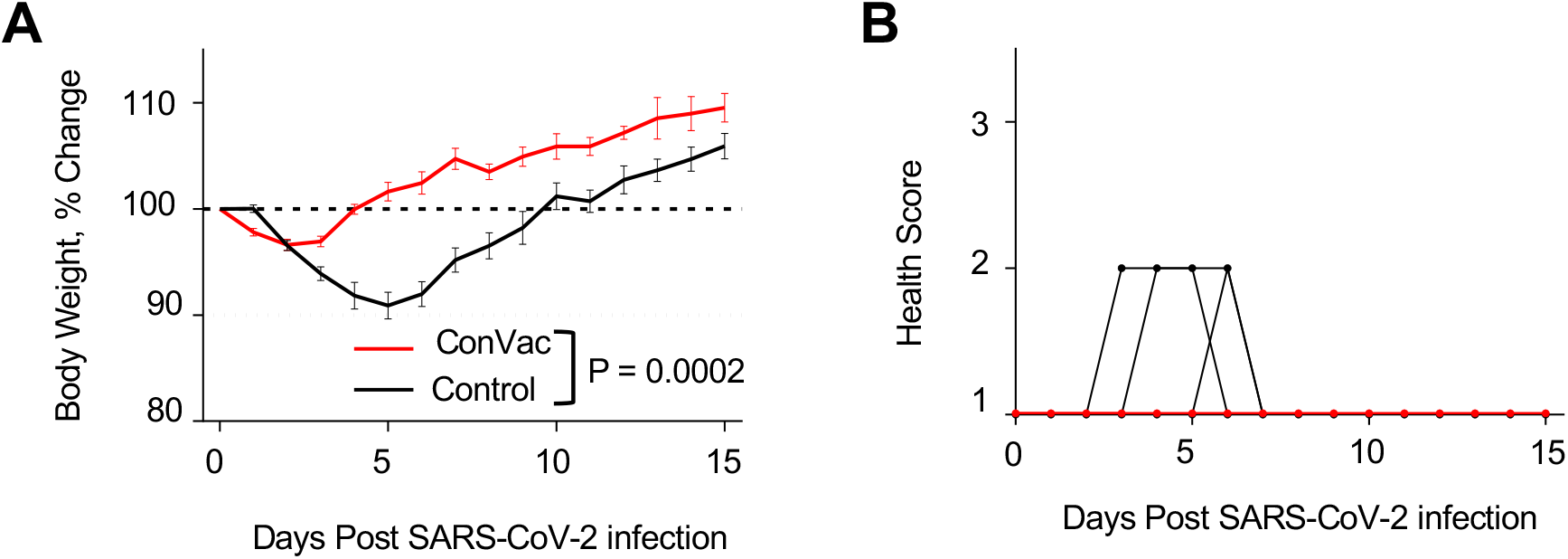
Hamster body weight change and health scores upon SARS-CoV-2 infection. Hamsters were vaccinated at day 0 and challenged intranasally with 10^5^ PFU SARS-CoV-2 at day 31. (A) % change in body weight and (B) individual disease score. VSV vaccine is shown in red and control group in black. N = 12 for control group (6 hamsters euthanized at day 3 post challenge) and N = 10 for ConVac group (5 hamsters euthanized at day 3 post challenge). Body weight P value determined by Wilcoxon test.

On days three and fifteen post challenge, half of the hamsters in each study group were euthanized, and lungs and nasal turbinates were harvested to determine viral loads by plaque reduction assay (Fig. 5A,B) and number of viral copies by RT-qPCR assay (Fig. 5C,D). In the control group three days post infection, high virus load was detected the lungs of all animals (Fig. 5A), ranging from 6.5×10^5^ PFU/g to 2.5×10^6^ PFU/g, and in the nasal turbinates of all but one animal ranging from 4.0×10^2^ PFU/g to 1.2×10^4^ PFU/g (Fig. 5B). In contrast, no virus was detected in the lungs of 4 out of 5 vaccinated hamsters, while the remaining one had the viral titer 645-fold less than in the control group. No virus was detected in the nasal turbinates of two vaccinated animals, while the three remaining animals displayed titers reduced by 57 fold compared to the control animals. On day 15, no SARS-CoV-2 was detected in lungs and nasal turbinates of both the control and the vaccinated hamsters (Fig. 5A,B).

**Figure 5.**
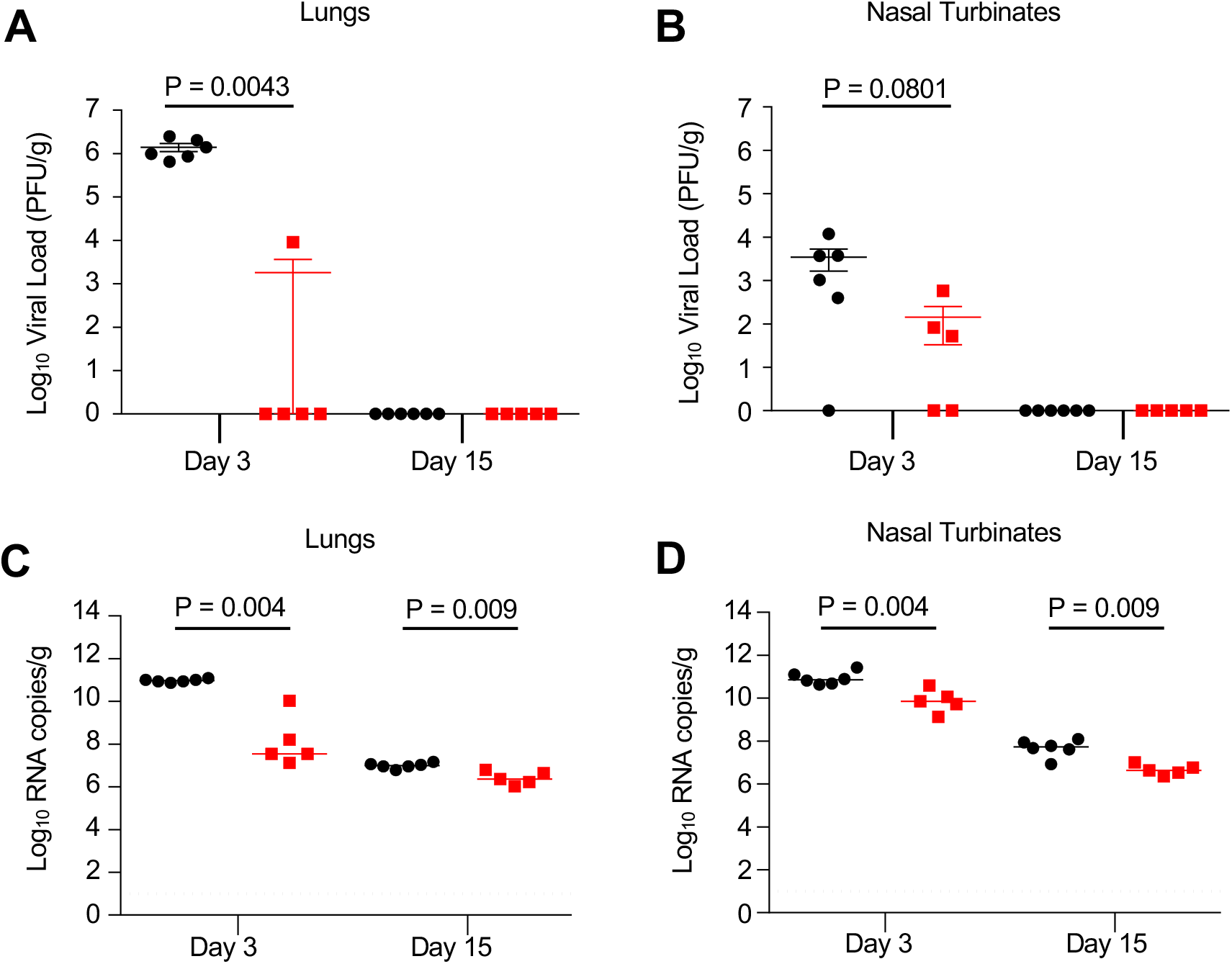
SARS-CoV-2 tissue viral load in hamsters. Hamsters were challenged intranasally with 10^5^ PFU SARS-CoV-2 and half of the animals in each group were euthanized at days 3 and 15 post challenge. Right lungs (A,C) and nasal turbinates (B, D) from each animal were homogenized in media and viral load were determined by plaque assays on Vero E6 cells (A,B) or by qRT-PCR (C,D). The limit of detection for the plaque assay was 70 PFU per lung and 35 PFU per nasal turbinate. The limit of detection for the qRT-PCR assay is indicated by the dotted line. The ConVac vaccine group is shown in red and the control group in black. For each timepoint, N = 6 for control group and N = 5 for ConVac vaccine group. P values determined by Mann-Whitney test.

RNA isolated from the lungs and nasal turbinate homogenates were assessed for the presence of viral RNA copies by RT-qPCR assay (Fig. 5C,D). In the control group, high virus RNA copies were detected in the lungs and nasal turbinates of animals with ~1×10^11^ RNA copies/g on day 3 post challenge, ~1×10^7^ RNA copies/g in the lung and ~6×10^7^ RNA copies/g in the nasal turbinates on day 15 post challenge. In contrast, significantly lower viral RNA copies were detected in the ConVac vaccinated animals with ~7×10^7^ RNA copies/g in the lungs and ~1×10^10^ RNA copies/g in the nasal turbinates on day 3 post challenge; ~3.6×10^6^ RNA copies/g in the lungs and ~5×10^6^ RNA copies/g in the nasal turbinates on day 15 post challenge. We assume that a portion of the detected viral RNA on day 3 is residual input challenge virus as it was below the detection limit of the plaque assay. In addition, the detected viral RNA on day 15 post challenge is not derived from viable viral particles since no plaques were detected.

### Vaccinated animals do not demonstrate enhanced lung pathology upon challenge, but still show some inflammation in the lungs

Pathology scores were determined in a blinded manner in lung sections of control and vaccinated animals on days 3 and 15 post challenge (Fig. 6,7). Regardless of the timepoint or the animal group, interstitial pneumonic changes were noted in all animals (Fig. 6, representative pathology pictures; and Fig. 7A,F, mean overall pathology scores). On day 3, the control group (Fig. 6A,B) showed widespread consolidation, inflammatory infiltration and obstruction of airways by inflammatory cells, while the ConVac vaccine group (Fig. 6C,D) demonstrated only moderate inflammatory changes with mononuclear cellular infiltration, septal thickening, and mild airway pathology. On day 15 (Fig. 6E,F), the control unvaccinated group displayed typical interstitial pneumonic changes consisting of small to large inflammatory foci, septal thickening and airway occlusion, with fewer inflammatory cells, while the ConVac vaccine group (Fig. 6G,H) displayed moderate interstitial pneumonia with septal thickening and reduced mononuclear cellular infiltrates in airways, but some hyperplasia was still present. These findings were confirmed by determining individual histopathology scores, which identified small to large inflammatory foci of moderate intensity involving large areas of lung sections on both days 3 and 15 (Fig. 7B-E,7G-J). Most animals had infiltration of large and small airways with inflammatory cells; cellular infiltrates consisted of mononuclear cells, including macrophages and lymphocytes. A moderate level of inter-alveolar septal thickening was a consistent feature in both groups (Fig. 7E,J). Perivascular cuffing was common and present in most sections. A semi-quantitative comparison of the control and vaccinated animals demonstrated no enhanced pathology in the vaccinated group (P>0.05 for all panels, Mann-Whitney test).

**Figure 6.**
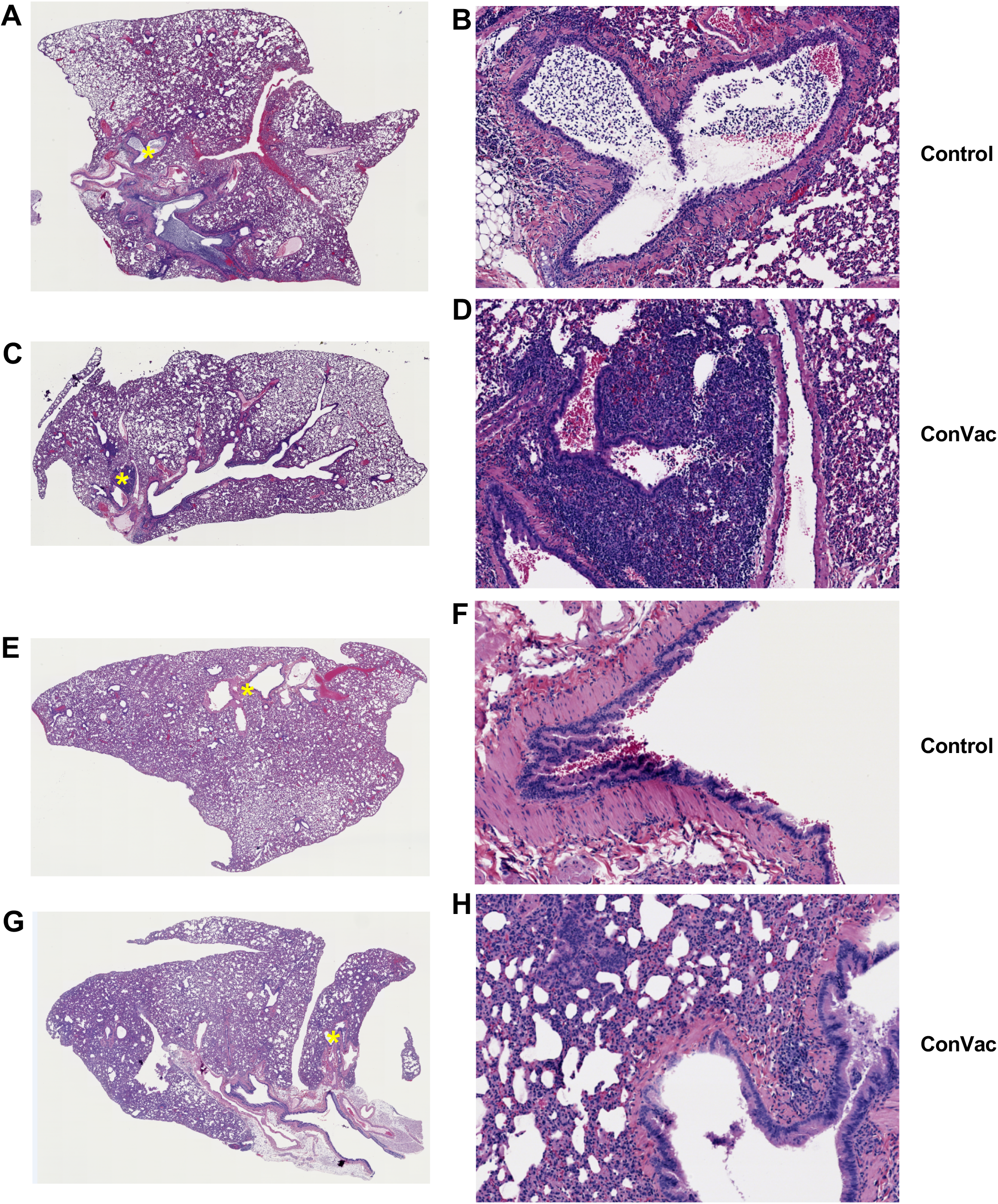
SARS-CoV2 Lung pathology. Representative histopathology of SARS-CoV-2 infection in control and vaccinated hamster lungs. Left panels: low magnification, right panels: higher magnification of selected regions, indicated by yellow asterisks on low magnification images. Magnified images are representative of specific pathological findings, not the extent or severity of pathology in a particular experimental group. A,B. Day 3 control: widespread consolidation and inflammatory infiltration is clearly visible, along with obstruction of airways by inflammatory cells. Higher magnification illustrates airway infiltration by mononuclear cells, septal thickening and airway occlusion. C,D. Day 3 ConVac vaccine: moderate inflammatory changes with mononuclear cellular infiltration, septal thickening, and mild airway pathology. Higher magnification illustrates focal infiltration by inflammatory cells. E,F. Day 15 control: typical interstitial pneumonic changes consisting small to large inflammatory foci, septal thickening and airway occlusion, with fewer inflammatory cells. Higher magnification illustrates epithelia hyperplasia and damage in airways. G,H. Day 15 ConVac vaccine: moderate interstitial pneumonia with septal thickening and mononuclear cellular infiltrates. Magnified image illustrates reduced cellular infiltration of airways, but hyperplasia is still present.

**Figure 7.**
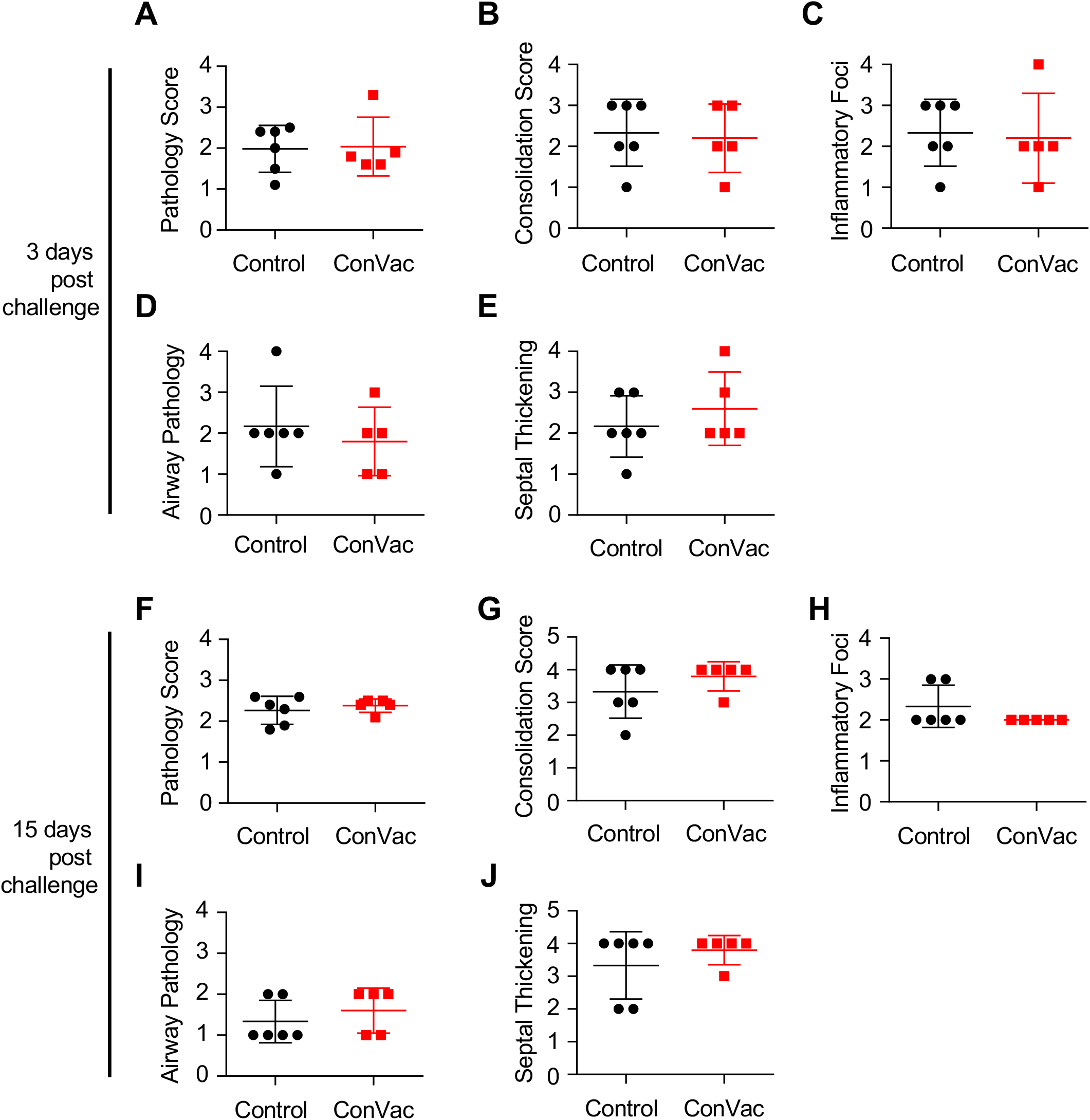
Comparative pathology scores for lungs from vaccinated and control hamsters post SARS-CoV-2 challenge. Scores at 3 days (A-E) and 15 days (F-J) post challenge. Scores for overall lung pathology (A, F), and individual criteria including consolidation or extent of inflammation (B,G), type inflammatory foci (C,H), airway pathology (D,I) and septal thickening (E,J) are displayed. The pathology scores (mean) were calculated based on the criteria described in Supplemental Table 1. The ConVac vaccine group is shown in red and the control group in black. Data represent mean ± SD, N = 6 for control group and N = 5 for ConVac vaccine group at each time point.

## DISCUSSION

The VSV vaccine platform has several features that make it a strong candidate for further development. These include the ability to stably express an inserted foreign open reading frame, the robust replication resulting in a high immunogenicity and the lack of preexisting immunity in the human population. VSV-vectored vaccine candidates have been developed against several human pathogens (Fathi et al., 2019) including Ebola virus; in 2019, the VSV-vectored vaccine Ervebo was approved by the European Medicines Agency and the FDA. Here we designed and developed a VSV-based SARS-CoV-2 vaccine ConVac and tested it for immunogenicity and efficacy in the hamster model of COVID-19.

The ConVac vaccine elicited a robust humoral response and protected hamsters from SARS-CoV-2 replication in the lower respiratory tract. A single dose of ConVac elicited strong SARS-CoV-2 S1 binding and virus neutralizing antibody titers. We note that two vaccinated animals were euthanized on days 10 and 11 due to a disease of unknown ethiology; however, we cannot completely exclude a residual neurovirulence of the vaccine construct at the high, 2 x 10^7^ FFU, vaccine dose used in our study. Viral loads determined by plaque assay in lung tissues were significantly reduced in ConVac-vaccinated animals compared to controls, with only one vaccinated hamster having detectable viral loads on day 3, suggesting that the ConVac vaccine prevents or strongly reduces SARS-CoV-2 replication in the lower respiratory tract. The hamster with the detectable viral load on day 3 had the lowest neutralizing antibody titer. Viral loads in the nasal turbinate tissues of vaccinated hamsters were reduced compared to control animals, but the difference was not statistically significant due to the limited number of animals per group and high variability between samples. Viral RNA levels determined by RT-qPCR confirmed these findings of a more robust protection in the lung compared to the nasal turbinates. These data suggest that ConVac elicited an immune response to effectively control the viral load.

Two recent studies used G gene-deleted VSV vaccine constructs expressing full-length SARS-CoV-2 S protein in mice and hamsters (Case et al., 2020; Yahalom-Ronen et al., 2020). In contrast to these two other studies, we used only the S1 subunit of the protein because it contains the necessary neutralizing epitopes with that in the receptor-binding domain being highly protective (Ho, 2020) and both prefusion and postfusion S2 conformations present potential drawbacks as vaccine constructs (Cai et al., 2020). Furthermore, unlike the S1 protein, the S2 subunit is more conserved between coronaviruses possibly leading to predominant S2-specific responses in humans previously infected with heterologous coronaviruses. The 50% mean neutralizing titer induced by our vaccine, 1:369, was higher than the titer of 1:173 elicited by the other VSV vaccine candidate assessed in hamsters by Yahalom-Ronen et al. (Yahalom-Ronen et al., 2020). When tested in mice, one of these vaccines elicited mean and median neutralizing titers greater than 1:5,000 after two vaccine doses but the median neutralization titer after one vaccine dose was only 1:59 (Case et al., 2020). The difference in the immunogenicity can also be explained by differences in growth kinetics and tissue tropism between a G-deleted VSV and a G-containing virus with the latter growing faster, having a broader tropism in vivo and targeting additional types of immune cells. Side-by-side comparisons would be needed to elucidate this further.

We used the hamster model of COVID-19, which does not require SARS-CoV-2 adaptation, exhibits high viral titers and severe pathologic changes in the lungs (Chan et al., 2020; Imai et al., 2020; Sia et al., 2020) and therefore seems to reproduce a severe form of COVID-19 observed in humans. Indeed, we confirmed that SARS-CoV-2 intranasal infection of hamsters results in viral replication in the upper and lower respiratory tract. In addition, lung histopathological damage was consistent with the changes described by others as this manuscript was in preparation (Chan et al., 2020; Imai et al., 2020; Sia et al., 2020). Therefore the golden Syrian hamster represents an excellent model of a severe COVID-19 suitable for vaccine testing.

Besides hamsters, several animal models have been developed for COVID-19. These include the mouse model with the mouse-adapted SARS-CoV-2 (Dinnon et al., 2020; Gu et al., 2020), transgenic mice expressing human angiotensin-converting enzyme 2 (ACE-2)(Bao et al., 2020); mice infected with a replication-defective Ad5 adenovirus expressing ACE-2 (Ad5-hACE2) transiently (Hassan et al., 2020) and the NHP model of COVID-19 (Munster et al., 2020). The mouse-adapted strains of SARS-CoV-2, despite reaching the titers of 10^6^ PFU/g or 10^8.3^ viral RNA copies per gram of lung tissue, caused only mild to moderate lung disease (Dinnon et al., 2020; Gu et al., 2020). In the Ad5-hACE2 transduced mice, SARS-CoV-2 replicated in the lungs reaching 10^6^ PFU/g and caused a lung pathology consistent with a severe pneumonia (Hassan et al., 2020). Transgenic mice expressing human ACE2 supported SARS-CoV-2 replication in the lungs (10^6.8^ copies/ml) and developed moderate interstitial pneumonia (Bao et al., 2020). In Rhesus macaques, SARS-CoV-2 replicated in the lungs reaching 10^5^-10^8^ copies/g but induced only mild to moderate interstitial pneumonia (Munster et al., 2020). Thus the levels of pathology induced by SARS-CoV-2 in the available animal models vary significantly from very mild to severe with the hamster representing a model of a severe disease, potentially explaining some residual lung pathology observed in the ConVac-vaccinated hamsters. Indeed, we propose that despite the lack of detectable viable SARS-CoV-2 by plaque assay, the observed pathologic changes in ConVac-vaccinated hamsters are likely to be related to the extremely high susceptibility of hamsters to SARS-CoV-2 on one hand and the very robust immune response to the vaccine on the other hand. Upon intranasal challenge, binding to cells of the epithelium of the respiratory tract and possibly some very limited replication, the virus is likely to be neutralized by the high levels of antibodies, resulting in the influx of Fc-bearing immune cells including lymphocytes and macrophages. Importantly, while some pathologic changes were observed in vaccinated animals, no enhancement compared to the control group was detected. Furthermore, testing of a vaccine in different animal models (mouse, hamster, ferret, NHP) with various levels of susceptibility to SARS-CoV-2 is likely to result in a more realistic prediction of the outcome of vaccination and subsequent infection in humans.

In conclusion, our data show that the VSV-vectored SARS-CoV-2 vaccine based only on the S1 subunit induces a high titer of serum neutralizing antibodies and protects hamsters from SARS-CoV-2 replication in the lower respiratory tract without causing an enhanced disease.

## MATERIALS AND METHODS

### Antibodies

Monoclonal antibody CR3022 was produced by transient transfection of 293F cells with cDNA expression plasmids obtained from BEI resources. The cells were transfected at a density of approximately 2×10^6^ cells/ml with equal amounts of each plasmid using Expifectamine reagent (Thermo Fisher) according to the instructions provided by the manufacturer. Supernatant was harvested after 4-5 days and antibody purified on Protein G Sepharose (Abcam, GE Healthcare) following standard protocols. Monoclonal antibodies I1 and I14 against the VSV glycoprotein and monoclonal antibody 23H12 against the VSV matrix protein were purified from hybridoma supernatant by affinity chromatography on Protein G sepharose. The antibodies were labeled with fluorophores using Dylight antibody labeling kits from Thermo Fisher. The following SARS-CoV-2 specific human monoclonal antibodies were kindly provided by Distributed Bio: DB_A03-09, 12; DB_B01-04, B07-10, 12; DB_C01-05, 07, 09, 10; DB_D01, 02; DB_E01-04, 06, 07; DB_F02-03.

### Generation of ConVac vaccine

Codon-optimized cDNA encoding the SARS-CoV-2 S1 domain fused to the C-terminal portion of the VSV glycoprotein was obtained from Genscript. The cDNA was cloned into the XhoI and NheI sites of a modified recombinant VSV vector containing an additional transcription start stop signal between the G and L genes (Wirblich et al., 2017). The recombinant virus was recovered on 293T cells as described previously, filtered through an 0.22 μm filter and used to inoculate Vero (CCL-81, ATCC) or human BEAS-2B lung cells (gift from R. Plemper, University of Georgia). The infected cells were cultured in serum-free VP-SFM or Optipro medium. Cell culture supernatant was harvested three days post inoculation, filtered through 0.45 um PES membrane filters and used for virus characterization and animal experiments.

### Characterization of the ConVac vaccine

Baby hamster kidney (BSR) cells were inoculated with ConVac or VSV control virus expressing GFP at an MOI of 0.2 PFU. The following day cells were lysed in detergent buffer (1% TritonX-100, 0.4% sodium deoxycholate, 150 mM NaCl, 20mM HEPES pH 7.0, 1 mM EDTA) supplemented with a protease inhibitor cocktail (Thermo Fisher). Protein concentration was determined by BCA assay (Thermo Fisher), and ten micrograms of total protein were resolved on 4-20% Tris-Glycine gels (Thermo Fisher). The proteins were transferred to nitrocellulose membrane and probed with polyclonal rabbit serum against SARS-CoV S1 domain (Thermofisher, Cat No. PA5-81798 and PA5-81795), monoclonal antibodies I1 (8G5F11) and I14 (IE9F) against the VSV glycoprotein and monoclonal antibody 23H12 against VSV matrix protein provided by Douglas Lyles (Wake Forest University). After incubation with HRP-conjugated anti-rabbit or anti-mouse IgG (Jackson Immunoresearch), the membrane was incubated with WestDura substrate (Thermo Fisher), and bands were visualized on a Fluochem M imager (Biotechne).

For immunofluorescence staining, Vero E6 cells were seeded on coverslips and infected at three different MOI ranging from 0.01 to 0.1 PFU. After 10 hours, the cells were fixed for 10 min in Cytofix/Cytoperm (Becton Dickinson) and permeabilized for 10 minutes in 0.5% Tween 20 in Dulbecco phosphate buffered saline (DPBS). After washing with DPBS the cells were incubated for 2 hours with monoclonal antibody CR3022 conjugated to Dylight 550 or a mixture of monoclonal antibodies I1 and I14 conjugated to Dylight 488. Coverslips were washed in DPBS and mounted using Prolong Glass (Thermo Fisher). Images were acquired on a Zeiss Axioskop40 microscope equipped with a ProgresCF digital camera.

To assess viral replication of ConVac *in vitro*, Vero (CCL-81) cells were seeded in 6-well plates and infected next day at 34°C at an MOI of 5 and 0.05 PFU. After 1.5 hours the inoculum was removed the cells were washed 3x in DMEM and fresh medium containing 2% FCS was added to each well. Supernatant was collected at various time points from 6 to 36 hours post infection. A ten-fold serial dilution of supernatant was prepared in DMEM medium in a 96-well plate. Diluted supernatant was added to Vero E6 cells seeded in 96-well plates the day before. After 24 hours incubation at 34°C, the cells were fixed with 80% acetone and stained with fluorescently labeled monoclonal antibody 23H12 against the VSV matrix protein.

### Viruses

The SARS-CoV-2 strain used in this study is the first U.S. isolate SARS-CoV-2 USA_WA1/2020 from the Washington Sate patient identified on January 22, 2020 (Harcourt et al., 2020). Passage 3 was obtained from the World Reference Center for Emerging Viruses and Arboviruses (WRCEVA) at UTMB. Virus stocks were propagated in Vero E6 cells. The challenge stock used in this study is passage 5. The recombinant SARS-CoV-2 expressing Neon Green protein (SARS-CoV-2-mNG) (Xie et al., 2020) used in the neutralization assay was developed by Dr. Pei-Yong Shi at UTMB. Virus stocks were propagated in Vero E6 cells and a passage 4 was used in this study.

### Animal studies

The studies were carried out in strict accordance with the recommendations described in the Guide for the Care and Use of Laboratory Animals of the National Research Council. UTMB is an AAALAC-accredited institution and all animal work was approved by the IACUC Committee of UTMB. All efforts were made to minimize animal suffering and all procedures involving potential pain were performed with the appropriate anesthetic or analgesic. The number of hamsters used was scientifically justified based on statistical analyses of virological and immunological outcomes.

### Pathogenicity of SARS-CoV-2 in hamsters

On day 0, seven week-old golden Syrian female hamsters (Envigo) were anesthetized with ketamine/xylazine, and 8 animals were exposed intransally to the targeted dose of 10^5^ PFU of SARS-CoV-2 in a volume of 100 μl, while 4 animals were mock-infected with 100 μl 1X DPBS. Animals were monitored daily for weight loss and signs of disease. Half of the animals in each group (4 SARS-CoV-2 infected and 2 mock-infected hamsters) were euthanized by CO_2_ inhalation for viral load determination 2 and 4 days post challenge.

### Vaccination and SARS-CoV-2 challenge

Seven week-old golden Syrian female hamsters (Envigo) were anesthetized with 5% isoflurane prior to immunization and blood collections and with ketamine/xylazine prior to the SARS-CoV-2 challenge. On day 0, the vaccine group (N=12 animals) was inoculated with 2×10^7^ FFU of ConVac in 100 μl injection volume via the intramuscular route (50 μl per hind leg), while the control group (N=12 animals) remained naive. Vena cava blood collections were performed one day prior to the immunization and on day 28 afterwards. On day 31, vaccinated and control animals were exposed intransally to the targeted dose of 10^5^ PFU of isolate SARS-CoV-2. Animals were monitored daily for weight loss and signs of disease. Half of the animals in each group (5 vaccinated and 6 control hamsters) were euthanized by overdose of injectable anesthetics 3 days post challenge for viral load determination. Remaining animals were euthanized 15 days post infection by overdose of ketamine/xylazine.

### Binding antibody response

To determine antibody responses to the S protein of SARS-CoV-2, an indirect ELISA was developed utilizing purified S1 protein. The soluble S1 protein was produced by transfecting 293T cells with a plasmid that expresses a secreted S1 ectodomain (aa 16-681) fused to the C-terminal HA tag. Purification of the HA-tagged protein from the supernatant of transfected cells was carried out as described previously (Kurup et al., 2015). Antibody responses to SARS-CoV-2 spike protein (S1) were measured by an indirect ELISA as described previously(Kurup et al., 2015). Briefly, wells were coated overnight at 4°C with 500 ng/mL of S1 recombinant protein. The secondary antibodies used in the ELISA are HRP-conjugated goat anti-syrian hamster IgG secondary antibody (Jackson immunoresearch, Cat# 107-035-142, 1:8000 in PBST) or mouse anti-hamster-IgG2/3-HRP (Southern Biotech, Cat# 1935-05, 1:8000 in PBST). Optical density was measured at 490 nm and 630 nm using an ELX800 plate reader (Biotek Instruments). Data were analyzed with GraphPad Prism (Version 8.4.3) using 4-parameter nonlinear regression.

### Neutralizing antibody response

Sera collected from animals were tested for neutralizing capabilities against SARS-CoV-2. Briefly, serum samples were heat-inacitvated (30 minutes at 56°C). 10-fold diluted sera were further diluted in a 2-fold serial fashion, and 60 μl of each serum dilution was mixed with 60 μl of SARS-CoV-2-mNG (200 PFU) (Xie et al., 2020). The serum/virus mixtures were incubated for 1 hr at 37°C. 100 μl of the serum/virus mixtures were then transferred to Vero E6 cell monolayers in flat-bottom 96-well plates and incubated for 2 days at 37°C. Virus fluorescence was measured with a Cytation Hybrid Multi-Mode reader at 488 nm (Biotek Instruments).

### Tissue processing and viral load determination

For the pathogenicity study, animals from each study group were euthanized on days 2 and 4 post challenge, and the lungs and nasal turbinates were harvested. For the vaccine study, animals were euthanized on days 3 and 15 post challenge, and the lungs and nasal turbinates were harvested. For both studies, right lungs and nasal turbinates were placed in L15 medium supplemented with 10% fetal bovine serum (Gibco) and Antibiotic-Antimycotic solution (Gibco), flash-frozen in dry ice and stored at −80°C until processing. Tissues were thawed and homogenized using the TissueLyser II system (Qiagen). Tissue homogenates were titrated on Vero E6 cell monolayers in 24-well plates to determine viral loads. Plates were incubated 3 days at 37°C before being fixed with 10% formalin and removed from containment. Plaques were visualized by immunostaining with 1 μg/mL cocktail of 37 SARS-CoV-2 specific human antibodies kindly provided by Distributed Bio. As the secondary antibody, HRP-labeled goat anti-human IgG (SeraCare) was used at dilution 1:500. Primary and secondary antibodies were diluted in 1X DPBS with 5% milk. Plaques were revealed by AEC substrate (enQuire Bioreagents).

### qRT-PCR

Tissue homogenates were mixed with TRIzol Reagent (Life Technologies) at a 1:5 volume ratio of homogenate to TRIzol. The RNA extraction protocol for biological fluids using TRIzol Reagent was followed until the phase separation step. The remaining RNA extraction was done using the PureLink RNA Mini Kit (Ambion). The quantity and quality (260/280 ratios) of RNA extracted was measured using NanoDrop (Thermo Fisher). SARS-CoV-2 nucleoprotein cDNA was generated from RNA from Bei Resources (NR-52285) by One-Step RT PCR (SuperScript IV, Thermo Fisher) with primers SARS COV-2 N IVT F1 (5’-GAATTCTAATACGACTCACTATAGGGGATGTCTGATAATGGACCC-3’) and SARS COV-2 N IVT R1 (5’-GCTAGCTTAGGCCTGAGTTGAGTCAGCACTGCT-3’). The SARS-CoV-2 N standards were generated by *in-vitro* transcription of the generated SARS-CoV-2 N cDNA using the MegaScript T7 Transcription kit (Invitrogen), followed by using the MEGAclear Transcription Clean-Up Kit. Aliquots of 2 ×10^10^ copies/μL were frozen at −80°C. Five microliters of RNA per sample were run in triplicate, using the primers 2019-nCoV_N2-F (5’-TTACAAACATTGGCCGCAAA-3’), 2019-nCoV_N2-R (5’-GCGCG ACATTCCGAAGAA-3’) and probe 2019-nCoV_N2-P-FAM (5’-ACAATTTGCCCCCAGCGCTTCAG-3’).

### Histopathology

Following euthanasia, necropsy was performed, gross lesions were noted, and left lungs were harvested in 10% formalin for histopathological assessment. After a 24-hour incubation at 4°C, lungs were transferred to fresh 10% formalin for an additional 48-hour incubation before removal from containment. Tissues were processed by standard histological procedures by the UTMB Anatomic Pathology Core, and 4 μm-thick sections were cut and stained with hematoxylin and eosin. Sections of lungs were examined for the extent of inflammation, type of inflammatory foci, and changes in alveoli/alveolar septa/airways/blood vessels in parallel with sections from uninfected or unvaccinated lungs. The blinded tissue sections were semi-quantitatively scored for pathological lesions using the criteria described in Supplemental Table 1. Examination was performed with an Olympus CX43 microscope at magnification 40X for general observation and 100X magnification for detailed observation. Each section was scored independently by two trained lab members and as scores were in agreement, only one set is presented.

### Statistical analyses

were performed with GraphPad Prism for Windows (version 6.07). *P*<0.05 was considered significant.

### Biocontainment work

Work with SARS-CoV-2 was performed in the BSL-3 and BSL-4 facilities of the Galveston National Laboratory according to approved standard operating protocols.

## ACKNOWLEDGEMENTS

We thank the World Reference Center for Emerging Viruses and Arboviruses (WRCEVA) at UTMB for propagating and providing the viral isolate SARS-CoV-2 USA_Wa1/2020 (GenBank accession number MN985325.1) used in this study. We thank the UTMB Animal Resource Center for the support of the animal study. We thank Distributed Bio for its kind gift of 37 SARS-CoV-specific monoclonal antibodies used for immunostaining.

## AUTHOR CONTRIBUTIONS

Conceptualization, M.J.S., A.B., D.C.M., D.K. and C.W.; Investigation, D.C.M., D.K., C.W., A.J.R., C.M., N.K., N.S., S.P., M.A.H and J.M.W., Writing - Original Draft, D.C.M., A.B. and C.W.; Writing - Review & Editing, D.C.M., C.W., A.B. and M.J.S.; Visualization, D.C.M., A.B., D.K., C.W., A.J.R. and S.P.; Resources, P.Y.S., A.B. and M.J.S.; Supervision, Project Administration and Funding Acquisition, A.B. and M.J.S.; All authors read and approved the manuscript.

## SUPPLEMENTAL INFORMATION

**Supplemental Table 1.** Criteria for histopathology scoring

