## Supplemental Table 1 for "A single dose of replication-competent VSV-vectored vaccine expressing SARS-CoV-2 S1 protects against virus replication in a hamster model of severe COVID-19"

**Supplemental Table 1.** Criteria for histopathology scoring

|  | **Scores🡪** | **0** | **1** | **2** | **3** | **4** |
| --- | --- | --- | --- | --- | --- | --- |
| **A** | Extent of inflammation (% tissue involved) | 0 | <10 | 10-30 | 30-60 | >60 |
| **B** | Inflammatory foci type | No inflammation | Patchy inflammatory foci, few (<2) | Patchy inflammatory foci, many (>2) | Large inflammatory foci, few (<2) | Large inflammatory foci, many (>2) |
| **C** | Alveolar septa | Thin and delicate | Thickened in <10% HPF | Thickened in <30% HPF | Thickened in <60% HPF | Thickened in >60% HPF |
| **D** | Airways | Clear; no cells | Few cells in airway | Moderate cells in airway | More cells in air way; Epithelial hyperplasia | Occlusion of air way/epithelial hyperplasia or desquamation |
| **E** | Alveoli/ perivascular cuff/blood vessels/ pleuritis/cell types | Clear; no inflammatory cells | Few cells. Few PMN or MNC | Moderate cells/ PVC/mild congestion/  mild pleuritis/mostly MNC | More cells/PVC/ more congestion and pleuritis/more MNC and PMN | Abundant cells/large PVC/severe congestion or pleuritis/mixed cells |

The criteria were adapted from Matute-Bello et al., 2011.

HPF – high power field (>10x); PMN – polymorphonuclear cells/heterophils; MNC – mononuclear cells including lymphocytes and macrophages; PVC – perivascular cuff.
